# Unveiling DNA Origami Interaction Dynamics on Living Cell Surfaces by Single Particle Tracking

**DOI:** 10.1101/2024.12.23.628980

**Authors:** Indra Van Zundert, Elena Spezzani, Roger R. Brillas, Lars Paffen, Angelina Yurchenko, Tom F. A. de Greef, Lorenzo Albertazzi, Alessandro Bertucci, Tania Patiño

## Abstract

Due to the unique spatial addressability of DNA origami, targeting ligands (e.g. aptamers or antibodies) can be specifically positioned onto the surface of the nanostructure, constituting an essential tool for studying ligand-receptor interactions at the cell surface. While the design and ligand incorporation into DNA origami nanostructures are well-established, the study of cell surface interaction dynamics is still in the explorative phase, where in depth fundamental understanding on the molecular interactions remains underexplored. This study uniquely captures real-time encounters between DNA origami and cells in-situ using single particle tracking (SPT). Here, we functionalized DNA nanorods (NRs) with antibodies or aptamers specific to the epidermal growth factor receptor (EGFR) and used them to target EGFR-overexpressing cancer cells. SPT data revealed that ligand coated NRs selectively bound to the receptors expressed in target cancer cells, while non-functionalized NRs only display negligible cell interactions. Furthermore, we explored the effect of ligand density on the DNA origami, which revealed that aptamer-decorated NRs exhibit non-linear binding characteristics, whereas this effect in antibody-decorated NRs was less pronounced. This study provides new mechanistic insights into the fundamental understanding of DNA origami behaviour at the cell interface, with unprecedented spatiotemporal resolution, aiding the rational design of ligand-targeted DNA origami for biomedical applications.

## Introduction

Cell surface interactions regulate numerous biological processes, including signal transduction^1^, immune responses^2,3^, cell fate^4^ and adhesion^5^, thus playing a pivotal role in maintaining tissue homeostasis and disease. A deep understanding on the underlying mechanisms of such interactions is essential for the development of new targeted therapeutics and diagnostic tools^6,7^. Recent advancements in nanotechnology have provided new methodologies to investigate cell receptor targeting. Due to their tuneable size and surface properties, nanoparticles allow for the functionalization of ligands with varied density and distribution, which can be used to modulate targeting efficiency and activation of cell surface receptors.^8,9^ For instance, by localizing multiple ligands onto a single nanostructure, multivalent interactions with cell receptors can lead to a higher selectivity in nanoparticle targeting^10^.

Despite these exciting advances, a complete control over ligand density and precise positioning onto the nanoparticle surface is far from trivial. Recently, DNA origami has emerged as a powerful and promising tool to tackle this challenge. By folding a long single-stranded DNA (ssDNA) scaffold using smaller ssDNA oligonucleotides (staples), DNA origami allows for the generation of highly programmable 1D, 2D and 3D nanostructures with a full control over size and shape.^11–18^ One of the most appealing features of this approach is that the DNA staple strands can be used as handles for the attachment of different biomolecules, such as antibodies, peptides or native ligands.^19,20^ Moreover, the DNA staples are located at known specific positions, offering unparalleled precision over the patterning of ligands. This unique programmability and versatility have shed light on critical aspects of cell receptor targeting, such as the influence of precise ligand spacing on targeting selectivity^21–23^ and cell response activation^24^, establishing DNA origami a promising tool for studying cell surface interactions. In fact, over the past decade, significant efforts have been made to ensure DNA origami biocompatibility^25–27^and stability^28–30^ in biological environments, broadening its application to a wide range of biological settings, both in vitro and in vivo.^31–40^ For instance, DNA origami structures have been designed to deliver chemotherapeutic drugs directly to tumour cells via active targeting, minimizing damage to healthy tissue.^33^ In gene therapy, DNA origami can be used to deliver functional nucleic acids, such as small interfering RNA (siRNA) or plasmids, to specific cells to treat acquired or genetic diseases.^34,41^ For vaccine development, antigens can be positioned in a highly ordered and repetitive fashion on DNA origami, thereby enhancing the immune response.^42^

So far, most studies have focused on the evaluation of DNA origami selectivity, cellular uptake, intracellular fate or therapeutic efficacy in target cell populations.^31–36^ With this approach, pivotal details regarding the molecular interactions and binding dynamics of DNA origami at the bio-interface remain underexplored. In fact, knowledge about the cellular engagement of targeted DNA origami at the single molecule and single cell level is limited, and studies on the dynamics of cell-origami interactions are scarce. Moreover, most of the literature focuses on ensemble and endpoint measurements, such as cell binding and uptake after a certain time.

To address these gaps in our understanding, we explored the initial interactions between DNA origami and the cell membrane, a critical first step for cell targeting and drug delivery applications. We perform an in-depth investigation of the molecular interaction dynamics in space and time via single particle tracking (SPT), a highly sensitive technique that allows for the monitoring of individual DNA origami trajectories overtime with unique spatiotemporal resolution. Using SPT, we evaluate DNA origami diffusion, binding kinetics and internalization dynamics, which can provide valuable insights into the mechanisms governing DNA-origami based cell targeting.

To do so, we used DNA nanorods (NRs) functionalized with antibodies or aptamers specific to the epidermal growth factor receptor (EGFR) as our model platform. Accordingly, MDA MB 468 breast cancer cells, which exhibit high EGFR expression^43,44^, were selected as the target cell line while Hek 293T cells as control due to their significantly lower EGFR expression^45^. Using SPT, we monitored the trajectories of individual NRs in the cellular environment over time. The collected tracking data were subjected to multi-parametric analysis, providing insights into the diffusive behavior of DNA origami, selectivity, receptor engagement and overall binding kinetics and dynamics of DNA-origami-cell interactions (Figure 1).

**Figure 1.**
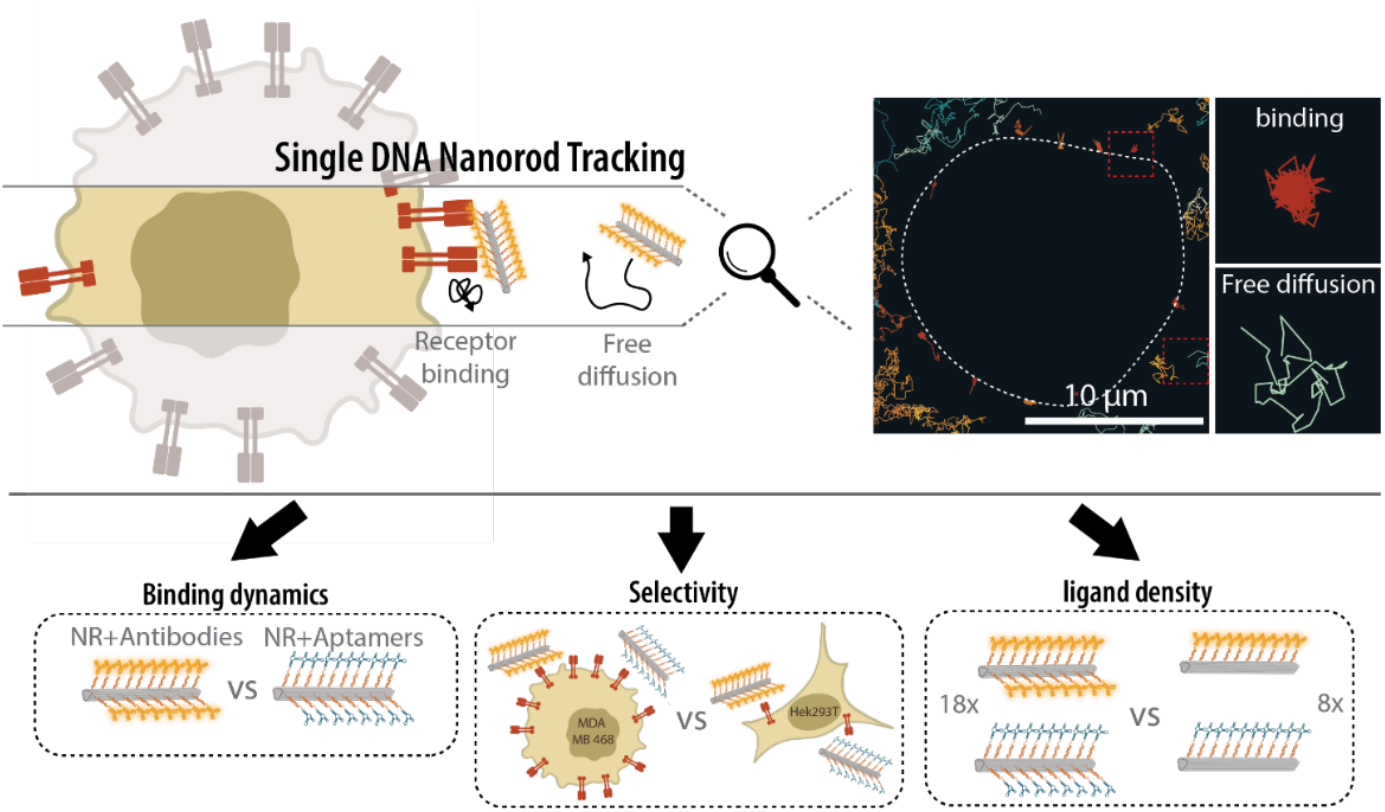
General approach for the single DNA-origami tracking on the cell surface. SPT was employed to monitor the trajectories of freely diffusing DNA origami nanorods and those bound to the cell membrane, where binding dynamics, selectivity and the effect of ligand density on receptor binding kinetics was evaluated.

## Results and discussion

To study the dynamics of DNA nanostructures at the cell surface, we employed an 18-helix bundle rod shaped DNA (15 x 150 nm) functionalized with either anti-EGFR antibodies or aptamers. The design of the NR allowed for the incorporation of a maximum of 18 antibodies or aptamers. Specifically, the NRs were adapted from Cremers et al.^23^ for accommodating 18 ssDNA handle sequences (Figure 2A) for antibody or aptamer hybridization and 6 ssDNA handles for Atto647N incorporation (Figure 2A). To incorporate the anti-EGFR antibody, we conjugated it to a ssDNA, complementary to the handle sequences, via click chemistry (see Method section). To verify the conjugation success, we performed native PAGE, where we observed an increased migration of the DNA-antibody conjugates towards the positive pole compared to the unconjugated antibodies. This can be explained by a higher negative charge provided by the DNA oligonucleotide, indicating a successful conjugation (Figure 2B). Moreover, we further confirmed these results by performing a SDS-PAGE, which showed an increase in molecular weight the DNA-antibody conjugate, which corresponds to the molecular weight of the conjugated ssDNA oligonucleotide (Figure S1). To specifically visualize the antibodies within the gel, we labelled them with Cyanine3 (Cy3) dye via NHS ester coupling. In the case of EGFR aptamer, we used a previously described design by Delcanale et al.^46^, which incorporated a ssDNA sequence complementary to the DNA origami handles. Both ss-DNA antibodies and aptamers were incubated with the DNA origami nanorods (Figure 2C) to allow for their hybridization with ssDNA handles. After that, the functionalized NRs (from now on, referred to as NR_18Ab and NR_18Apt, respectively), were analyzed by agarose gel electrophoresis. As expected, we observed a lower migration of both antibody- and aptamer-conjugated NRs compared to non-functionalized NRs, this effect being higher in the case of NR_18Ab. We attribute this difference to the higher molecular weight of antibodies compared to aptamers. We did not observe any band corresponding to non-functionalized NRs in the NR_18Ab or NR_18Apt lanes, indicating that the majority of NRs were successfully functionalized with their respective ligands. Lastly, functionalization was further confirmed by Atomic Force Microscopy (AFM) (Figure 2D).

**Figure 2.**
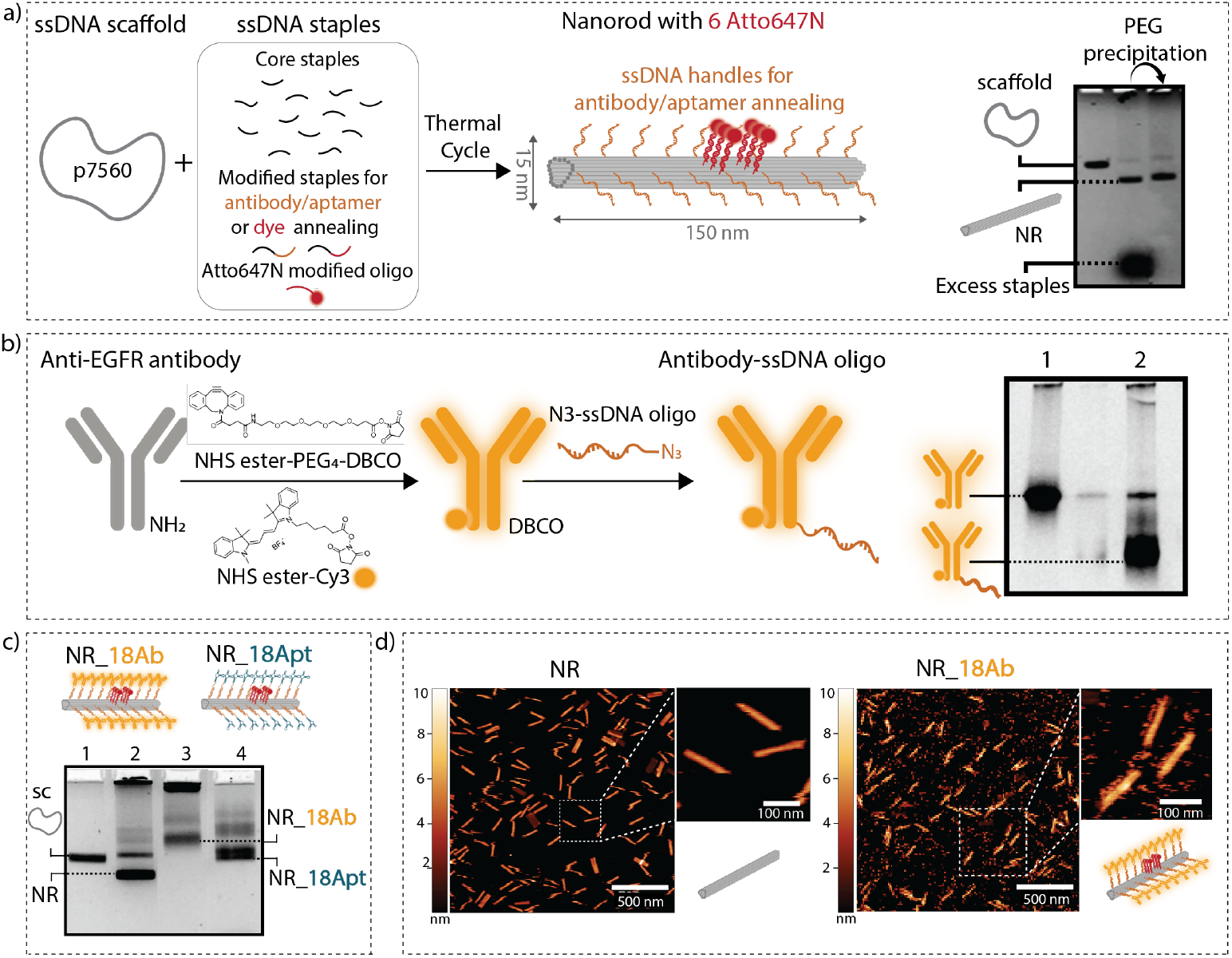
Fabrication and characterization of antibody- and aptamer-conjugated DNA nanorods. **a)** Schematic on the DNA origami self-assembly approach, combining the p7560 viral scaffold, the staple strands (core staples, 18 staples with extended handles for antibody/aptamer functionalization and 6 staples with extended handles for fluorescent dye functionalization). Folding and purification of the NR via poly-ethylene glycol precipitation was confirmed by agarose gel electrophoresis. **b)** Schematic representation of anti-EGFR conjugation approach to a ssDNA oligonucleotide, via click chemistry, and to Cy3, via NHS-ester. A native PAGE shows further migration of the oligo conjugated antibody towards the positive pole (lane 2). **c)** Conjugation of the anti-EGFR antibodies or aptamers to the NRs. The agarose gel shows differences the migration height of the different NRs, confirming their correct functionalization. Legend: Sc=Scaffold, NR=Folded DNA origami nanorod, NR_18Ab= NRs functionalized with anti-EGFR antibody, and NR_18Apt= NR functionalized with anti-EGFR aptamers. **d)** Atomic force microscopy images of the non-functionalized NRs (upper panel) and the NRs coated with 18 antibodies (lower panel).

Prior to studying DNA origami-cell interactions, we performed a fluorescence immunostaining of the EGFR in MDA MD 468 cells, which confirmed the expression of the receptor (Figure S2). Then, we proceeded to analyze the DNA origami interactions with the cell surface under an ONI microscope. For this, MDA MD 468 cells were cultured on glass bottom dishes overnight and different NR designs (non-functionalized NRs, NR_18Ab or NR_18Apt) were administered to the cells directly at the microscope at a final concentration of 1 nM. To visualize the NRs, we imaged the Atto647N fluorophores on their surface (Figure 1a). A minimum of 5 movies (10ms exposure time, 100 frames per second) were recorded at 10, 30 and 60 minutes after administration of the NRs. From the movies, the tracks of single NRs could be analyzed (tracking parameters are specified in the Methods section). Figure 3A shows representative images of non-functionalized NRs, NR_18Abs or NR_18 Apt trajectories at 30 min of incubation with cells (the trajectories of the other time points can be found in Figure S3). Cell contours (represented as dotted lines) were extracted from bright field images. With regard to non-functionalized NRs, we observed that most of the trajectories were characteristic of Brownian motion. From the trajectories, we could calculate the diffusion coefficients (Figure 3B). We observed a typical gaussian distribution of the diffusion coefficients for non-functionalized NRs, with an average diffusion coefficient of around 2.5 µm^2^/s at all three time points. By contrast, in both antibody- and aptamer-conjugated NRs, we observed two clear and distinct populations. On one hand, we observed a population that displayed the same diffusion behaviour as the non-functionalized NRs, with an average diffusion coefficient between 2 and 3 µm^2^/s, indicating Brownian motion (Figure 3B, blue shadow). On the other hand, we observed a population with a particularly low diffusion coefficient, with values between 0 and 1 µm^2^/s, (Figure 3B, yellow shadow). We attribute the low diffusion of this second population to cell surface binding. Indeed, Figure 3a shows that such low diffusive trajectories (red tracks) were always found in the vicinity of the cell membrane, indicating the presence of binding events between the antibody/aptamers on the NR and the cell membrane. In order to obtain further insights into NR motion dynamics, we represented the mean squared displacement (MSD) over time for the two distinct populations (i.e. high-diffusion, D > 1, and low-diffusion, D ≤ 1, populations). The MSD is a measure of the average distance a particle travels over time, often used to analyse the type of motion of a particle, which can be Brownian (linear MSD), ballistic (exponential MSD) or confined (plateau). For all the DNA origami designs within the high diffusion population, we observed linear MSDs, indicating Brownian motion. In the case of both aptamer and antibody functionalized NRs, the population with D ≤ 1, we observed a clear decrease of the MSD slope overtime, indicating confinement and reduction of the diffusivity of the particles, which we attribute to cell receptor binding (Figure S4).

**Figure 3:**
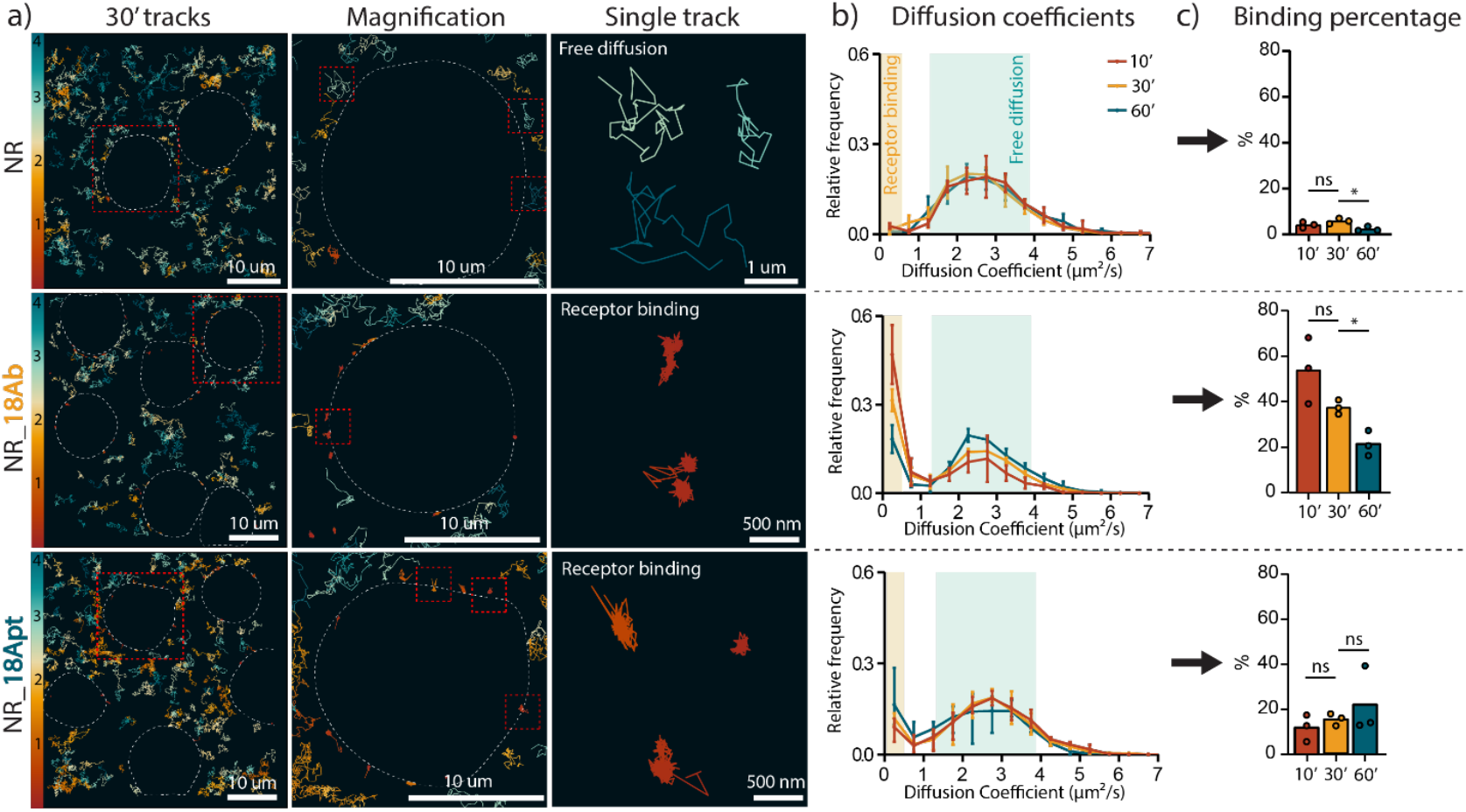
Characterization of NR diffusion and cell binding by SPT. **a)** Representative images of single particle tracking movies at 30 minutes after NR administration, showing single NR trajectories. Cell contours are indicated by dotted lines. Color bar indicates the diffusion coefficient (ranging from 0 to 4 *µ*m^2^/s). **b)** Histograms of the diffusion coefficients obtained from the NR trajectories at different timepoints 10 min (red), 30 min (yellow) and 60 min (blue) after NR addition to the cells. Cell binding events (NRs that have a diffusion coefficient less or equal to 1) are highlighted in yellow whereas NRs freely diffusing in the solution surrounding the cells are highlighted in light blue. **c)** Percentage of NR binding to the cells at different timepoints. n = 3 biological replicates (per biological replicate, the trajectories of 5 different movies were combined, i.e. 5 technical replicates); Results are shown as the mean i-standard error of the mean, where * indicates significant differences between groups with p ≤ 0.05.

Next, we calculated the average percentage of NRs binding to the receptor at different time points (10, 30 and 60 min, Figure 3C) by dividing the number of receptor-binding trajectories (i.e., the number of NRs with a diffusion coefficient ≤ 1) by the total number of trajectories. Non-functionalized NRs exhibited minimal receptor binding (under 3%), whereas both NRs_18Ab and NRs_18Apt began to interact with the receptor within 10 minutes, with a binding percentage of 54% and 12%, respectively. Although NRs_18Ab demonstrated higher initial binding (54% at 10 minutes), this percentage declined to 21% over 1 hour. By contrast, NRs_18Apt showed a lower initial binding (12% at 10 minutes) but increased marginally over time (22%, although this increase was not statistically significant). It is noteworthy that within the time frame of our experiment (60 min), NR internalization into the cell was scarce. To determine whether cell-bound NRs were eventually internalized at longer times of incubation, we evaluated cellular uptake at 24 hours post NR administration using confocal microscopy. Results showed that both NRs_18Ab and NRs_18Apt were internalized by MDA-MD-468 cells, whereas non-functionalized NRs showed lower cellular uptake (Figure S5).

After confirming that both antibody- and aptamer-conjugated NRs were able to interact with the EGFR at the cell surface, we examined the binding of NRs to a low-EGFR-expressing cell line (Hek 293T cells) to assess the impact of receptor density and origami selectivity (Figure 4A). To confirm the difference in EGFR expression in the two selected cell lines, an immunostaining was performed (Figure S2), which showed minimal fluorescence signal, confirming low EGFR expression. To investigate the differences in cell interactions between cells with high and lower EGFR expression, we conducted a comparative analysis using single particle tracking (Figure 4A). First, we focused on the analysis of NR trajectories. Figure 4B shows representative images of both cell types incubated with NR_18Ab during 60 min. We observed a higher number of ligand-functionalized NRs in MDA MD 468 cells compared to Hek 293T. Moreover, binding events of targeted NRs with MDA MD 468 cells were characterized by extremely long trajectories (often longer than 500 steps). By contrast, trajectories of NRs in the Hek 293T surface were significantly shorter. The length of a particle trajectory can be influenced by multiple factors. For instance, a trajectory could terminate because the respective NR moves out of the focal plane of the microscope (possibly while dissociating from the receptor after binding) or because the dye molecules on the NR undergo photobleaching, impeding the visualization of the DNA origami. Since the contribution of both moving out of focal plane and bleaching effects can be considered equal for all the NR designs, we explored the analysis of track length as a measure of the binding behaviour of NRs to the cell surface. To explore the differences in track lengths for different NRs design and different cell types, we analyzed the number of steps on the tracks of single NRs. Figure 4C shows the number of steps for each NR trajectory, plotted against their diffusion coefficient. We restricted the analysis to NRs which had a diffusion coefficient D ≤ 1, since these were considered as cell binding NRs. In accordance with the qualitative observations in Figure 4B, Figure 4C confirmed that the trajectories of antibody and aptamer conjugated NRs were significantly longer in the MDA MD 468 cells than in Hek 293T cells, with a p-value of 0.016 and 0.0002, respectively. Despite the absence of targeting moieties in the non-functionalized NRs, a small number of binding events could still be detected in both cell lines (Figure 4C, graph 1). Since the duration of these NR-cell interactions was rather short and no significant difference was found between the two cell types, these binding events were categorized as non-specific NR-cell interactions. Based on the empirical data, we found that those non-specific binding trajectories are typically shorter than 100 steps (which corresponds to 1 second bound time). This type of non-specific binding could also occur for the antibody and aptamer conjugated NRs. Therefore, to evaluate the selectivity of the targeted NRs, we considered only the specific binding events, defined as those that lasted longer than 100 steps (1 second). Figure 4D shows the percentage of specific binding events (trajectories with D ≤ 1 and a track length > 100 steps) for the different NR designs and cell types. Both antibody and aptamer decorated NRs (yellow and blue, respectively), display significantly higher specific binding to MDA MD 468 cells than to Hek 293T cells, demonstrating that ligand decorated NRs are highly selective towards the EGFR-overexpressing cell line. There were no significant differences found overtime (10, 30 or 60 minutes after NR addition), suggesting that specific interactions were stable in time.

**Figure 4:**
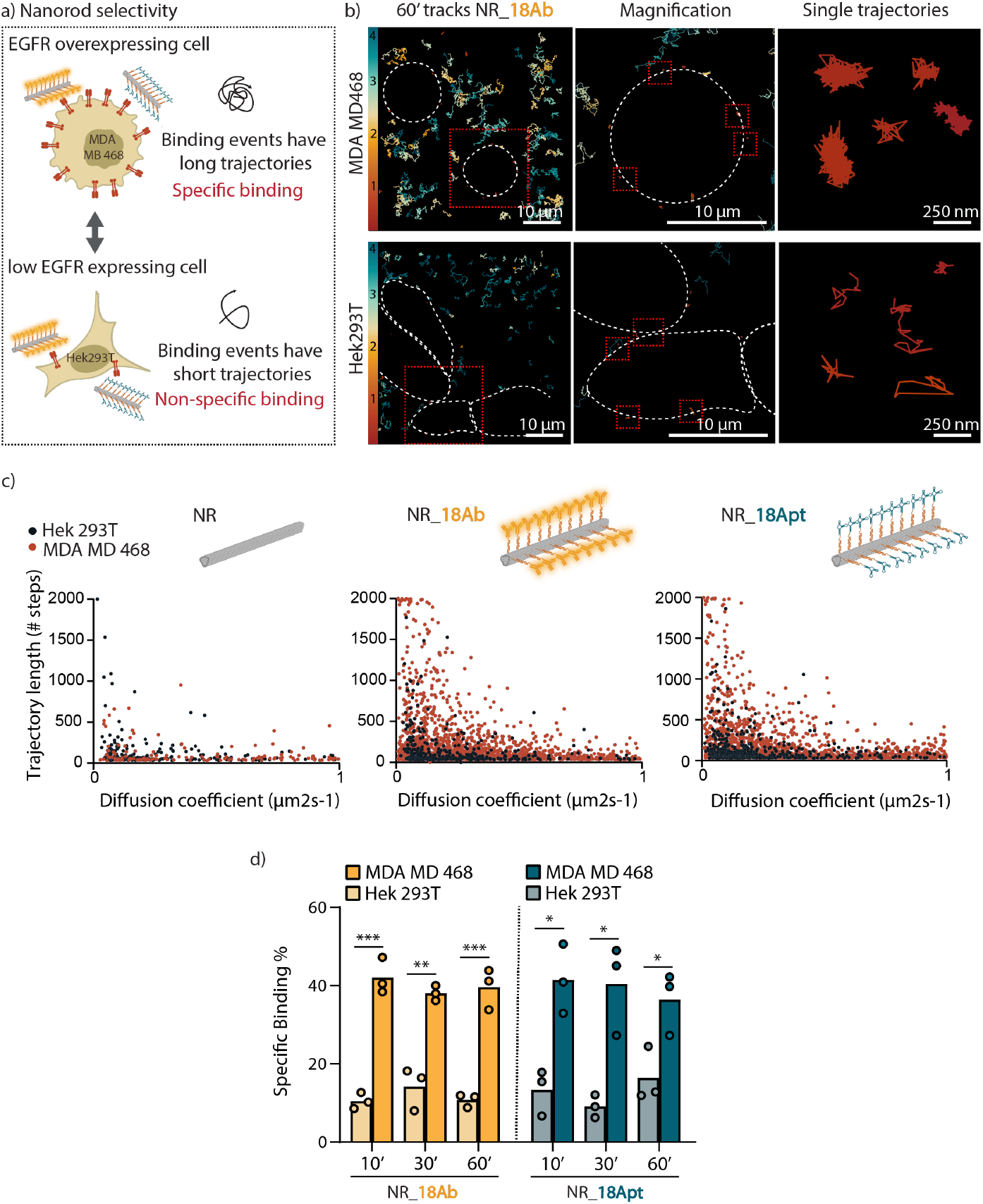
Selectivity of antibody and aptamer conjugated NRs. **a)** Schematic representation of the experimental findings on NR selectivity: the differential binding profile between the targeted cell line (MDA MD 468) and the non-targeted cell line (Hek 293T) with the targeted NRs (NRs_18Ab and NRs_18Apt). For the MDA MD 468 cells, binding events with the targeted NRs are characterized by very long trajectories, indicating specific binding, while the binding trajectories with the Hek 293T are typically much shorter, indicating non-specific binding. **b)** Representative image of NRs_18Ab trajectories at 60 minutes after their incubation with MDA MD 468 and Hek 293T cells. Cell contours are indicated by the dotted line. Color bar indicates the diffusion coefficients (ranging from 0 to 4 µm^2^/s). **c)** Scatter plot of all the binding events for non-functionalized NRs, NR_18Ab and NR_18Apt (i.e. all the trajectories with D ≤ 1), plotted against their respective trajectory length (y-axis). **d)** Bar plot displaying the differential specific binding percentage between MDA MD 468 and Hek 293T for the different NR designs (NR_18Ab and NR_18Apt) at 3 distinct time points (10 min, 30 min and 60 min). Results are shown as the mean +-standard error of the mean. n = 3 biological replicates (per biological replicate, the trajectories of 5 different movies were combined, i.e. 5 technical replicates) *: significant difference between groups with p ≤ 0.05, ** : significant difference between groups with p ≤ 0.01, *** : significant difference between groups with p ≤ 0.001.

Next, we evaluated the kinetic parameters characterizing the binding between the ligand coated NRs and the EGFR receptor, focusing on the effect of multivalency. For this, we engineered two versions of antibody- and aptamer-functionalized NRs, with either 8 or 18 ligands per NR (characterized in Figure S6). These configurations helped us to investigate the potential effects of multivalency in two distinct cellular contexts: high EGFR expression (MDA MB 468 cells) and low EGFR expression (Hek 293T cells). Multivalency refers to the ability of a single NR to engage with multiple receptors simultaneously by multiple ligands. In principle, multivalent interactions increase binding strength because a single particle can form several connections with the cell surface, making it more likely to remain bound even if one ligand-receptor interaction dissociates. These collective interactions often lead to stronger overall affinity (estimated by the dissociation constant, K_D_) than what would be expected from simply adding up the individual binding affinities of each ligand. We hypothesized that in cells with high receptor expression, multivalent interactions would enhance binding affinity beyond what is predicted by a linear increase in receptor density. This enhanced affinity and selectivity due to multivalency is often referred to as “super-selectivity”.^10,47,48^ By analysing the trajectories of single NR binding we can extract valuable information about two key kinetic determinants: the rate constant for ligand-receptor association (k_on_) and the rate constant for dissociation of the ligand-receptor complex (k_off_). A schematic representation of this concept is depicted in Figure 5A. First, we investigated the kinetics of binding events by analyzing the total number of specific interactions occurring on the cell surface (i.e. tracks with D ≤1 and a trajectory length > 100 steps). According to Equation 1 (Eq. 1 in the methods section), the number of binding events is directly proportional to the association rate constant (k_on_), the concentration of NRs (c_i_) in solution, and the number of receptors on the membrane (n). Since the receptor density (within the same cell line) and NR concentration are assumed constant across experiments, differences in the number of binding events reflect changes in k_on_. Here, we analyzed the total number of binding events to MDA MD 468 and Hek 293T cells for the different NR configurations (Figure 5B). It is worth nothing that, since no significant differences in specific binding were detected previously among the different time points (Figure 4D), data from the different time points were combined for this analysis. Overall, we observed that 18-ligand NRs exhibited a higher observed k_on_ compared to the 8-ligand NRs. This increase was even more pronounced for the aptamer-conjugated NRs, which showed a non-linear binding increase (approximately a 4-fold increase) for both cell lines, compared to a 1.3-1.4-fold increase for the antibody-functionalized NRs. The difference in the extent of binding enhancement between aptamers and antibodies can be attributed to several factors. One possibility is that antibody binding may have reached a saturation point where further binding is limited by diffusion or steric accessibility to the binding sites. However, in Hek 293T cells, where receptor density is lower, the relatively small 1.1x increase in binding for antibody-decorated NRs from 8 to 18 ligands suggests that the binding saturation is unlikely. Instead, the reduced accessibility of receptors, potentially due to the larger size and bulkiness of antibodies compared to aptamers, may be a limiting factor.^20,49,50^ Moreover, selectivity and multivalency are more enhanced in weak interactions, and therefore our results could also be explained by the fact that aptamers have a lower affinity than antibodies and therefore their binding was significantly enhanced when we increased the amount of ligands in the NR structure.^46^

**Figure 5:**
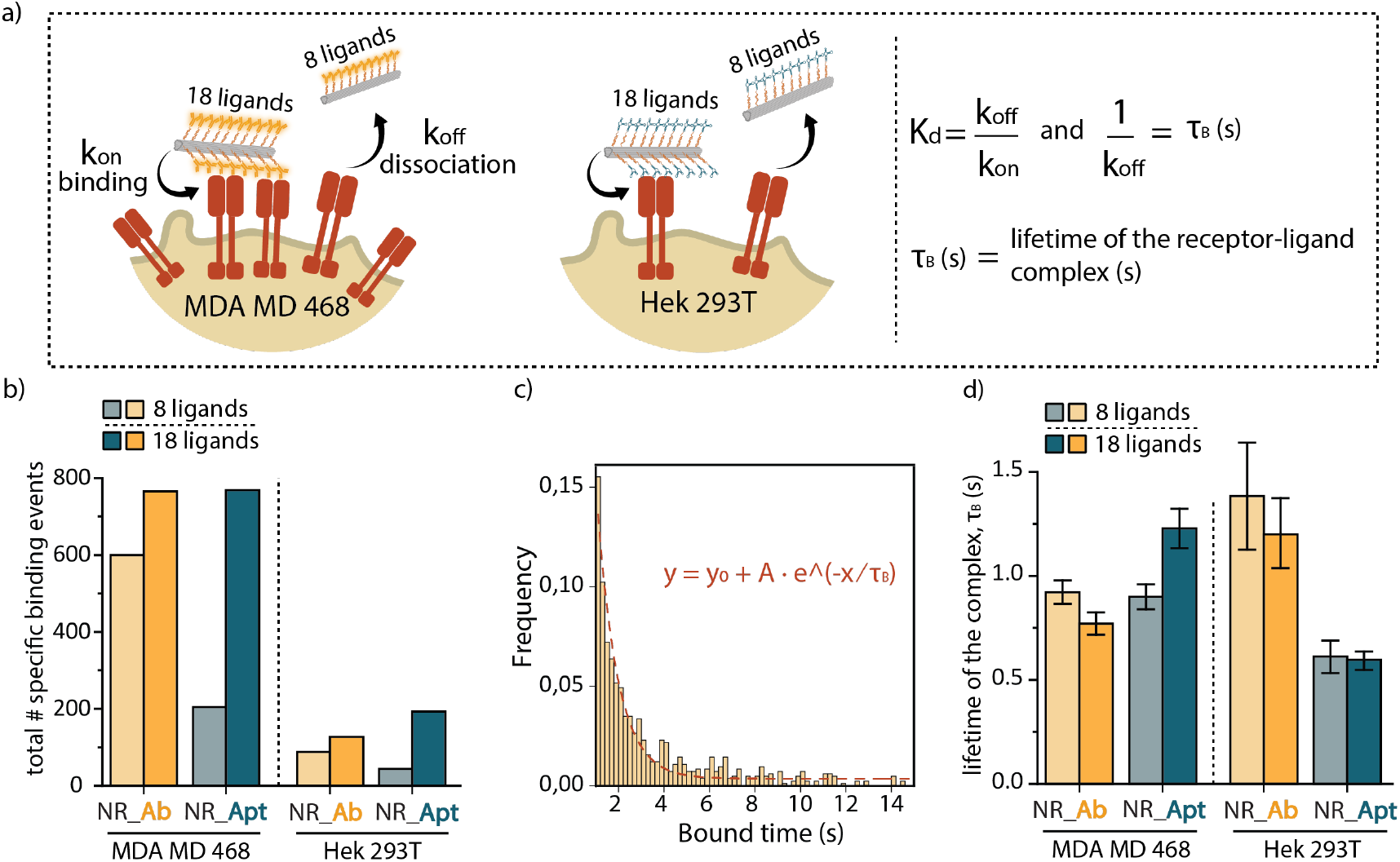
Binding kinetics of antibody and aptamer coated NRs. **a)** Scheme of the binding kinetics between targeted NRs (NR_18Ab and NR_18Apt) and cells (left panel). The corresponding formula of the binding kinetics is displayed in the right panel. **b)** Comparison of the total number of specific binding events between NRs with 8 or 18 binding ligands (antibody or aptamer) in both MDA MD 468 and Hek 293T cells. The amount of binding events can be directly related to k_on_. **c)** Example of the exponential decay fitting for the binding time of NR functionalized with 18 antibodies in MDA MD 468 cells, where dotted line represents the fitted courve and the corresponding equation is displayed. **d)** τ_B_ values obtained via an exponential decay fitting of all the binding events for each NR design and cell type. This value is inversely proportional to k_off_. A comparison was made between NRs with 8 or 18 binding ligands (antibody or aptamer) in both MDA MD 468 or Hek 293T. Results are shown as mean fitted value, where error bars represent the standard error of the fitting for each NR design in the different cell lines.

We next explored how different NR designs affect k_off_, which indicates how quickly the nanorods dissociate from the receptors. To quantify this, we fitted the distribution of binding event durations (i.e., the time the NRs remain bound to the receptors) with an exponential decay function to obtain τ_B_, which is inversely proportional to k_off_ (Eq. 2). An example of this fitting is shown in Figure 5C, where the bound time of NRs functionalized with 18 antibodies in MDA MD 468 cells was fitted. From these fittings, the τ_B_ value was obtained, which represents the average bound time and reflects receptor-NR complex stability. To compare the τ_B_ values among the different NR designs, they were plotted in Figure 5D. In the antibody-decorated NRs, the receptor binding time did not increase as the number of ligands increased. This result is surprising, as more ligands would typically be expected to promote multivalent interactions, thus prolonging binding duration. However, increasing the number of antibodies may introduce steric hindrance, limiting accessibility to available receptors. Moreover, the affinity of antibodies has been shown to change upon conjugation to a DNA origami platform, which could also explain our observations.^23^ This counterintuitive finding underscores the importance of k_off_ as a crucial design parameter for developing targeted DNA origami nanostructures. In contrast, the aptamer-coated NRs exhibit behaviour in line with our expectations. In MDA MB 468 cells, where EGFR density is high, the receptor binding time increases as the number of ligands increases, suggesting stronger multivalent interactions. Conversely, in Hek 293T cells, where receptor density is low, the binding duration remains unchanged, indicating that interactions are primarily monovalent (i.e., one aptamer binding to one receptor). This selective behavior of the aptamer-DNA origami design is in line with the super-selectivity principle, in which multivalent interactions enhance binding to cells with high receptor expression, a crucial feature for creating an efficient and specific targeting nanosystem.

## Conclusion

The exploration of DNA origami as a versatile platform for targeted drug delivery and cell response modulation is a rapidly evolving field, capitalizing on the structural precision and controllability of these DNA nanostructures. Recent research has produced various sophisticated DNA origami-based delivery vehicles. So far, most studies have focused predominantly on the design of the origami structures and end-point cellular uptake, and delivery efficiency analysis. A key challenge remains in understanding the molecular interactions between DNA origami and living cells. Here, we aimed at characterizing such interactions in real time, at single-particle and single-cell resolution, to investigate the dynamics and kinetics of DNA origami targeting *in situ*. We performed a quantitative analysis of the diffusive behaviour of DNA origami, as well as a multiparametric analysis of the molecular binding interactions in the native cellular environment, offering new insights into the molecular mechanisms underlying DNA-origami cell interactions. Our results demonstrate that both antibody- and aptamer-functionalized NRs can effectively engage with the EGFR, this engagement being selective towards EGFR overexpressing MDA-MB-468 cells. The selectivity of our ligand coated NRs was further confirmed through comparisons with low-EGFR-expressing Hek 293T cells. Thanks to the use of SPT, we could quantify not only the amount of NR binding to cells but also the duration of cell-receptor interactions with high precision, where both antibody- and aptamer-functionalized NRs displayed much longer binding durations compared to non-functionalized NRs. Moreover, this technique allowed to perform a systematic analysis of binding kinetic parameters, including the association rate constant (k_on_) and dissociation rate constant (k_off_), which provide highly valuable information about the strength and duration of ligand-functionalized DNA origami. Interestingly, we have observed that aptamer-coated NRs show non-linear binding behaviour in high-receptor density environments (e.g., increased binding and prolonged binding times with an increasing number of ligands), in line with the concept of super-selectivity. By contrast, for antibody-coated NRs, the addition of more antibodies to a NR did not show higher binding capacity, which is counterintuitive. Our results suggest that aptamers, due to their lower affinity, smaller size and better accessibility compared to antibodies, may be more effective in promoting multivalent interactions. This unexpected behavior highlights the complexities introduced by factors such as steric hindrance and receptor accessibility, suggesting that increasing the number of ligands does not always enhance receptor engagement and may even reduce DNA origami binding stability under certain conditions.

In summary, in this study we elucidate the mechanisms governing DNA-origami cell surface interactions, shedding light into the factors that modulate binding dynamics and kinetics. Our findings hold potential for advancing the design of DNA nanostructures for biomedical applications, including targeted drug delivery, biosensing and therapeutic applications.

## Acknowledgements

This work was supported by the Irene Curie Fellowship and the ICMS (T.P.). A.B. and E.S. acknowledge financial support under the National Recovery and Resilience Plan (NRRP, 2022FPYZ2N) and the COMP-R Initiative (MUR 2023-2027). I.V.Z. thanks the Flanders Research Foundation (FWO) (12A6N25N).

## Author contributions

The manuscript was written through contributions of all authors. / All authors have given approval to the final version of the manuscript. / **†** These authors contributed equally

## Materials and methods

### Materials

P7560 DNA scaffold (Tilibit nanosystems, Germany), Staples and modified staples (Integrated DNA technologies Europe), PEG8000 (Sigma-Aldrich, USA), InVivoMab anti-human EGFR antibody (Bio X cell, USA), NHS ester-(PEG)_4_-DBCO (Vector Labs, USA), Cyanine3 NHS ester (Lumiprobe, USA), Amicon 10 and 100 kDa weight cut-off (Sigma-Aldrich, USA), mini-PROTEAN TGX precast gels (BIO-RAD Laboratories Inc, USA)

### Nanorod Folding

The assembly of the NRs was performed by combining 20 nM of the viral scaffold (p7560) with 100 nM of each staple strand (5x excess of the staple strands), 12 mM MgCl_2_, 25 mM NaCl, 5 mM Tris (pH 8.5), and 1 mM EDTA. For fluorescence labeling, six staple strands were modified with a handle sequence complementary to an oligonucleotide modified with ATTO647N, which were added to the mixture at a final concentration of 100 nM. To enable surface decoration of the NR, 18 extended staple strands were incorporated at specific positions. The folding process commenced with an initial rapid heat denaturation step, followed by a cooling phase from 80°C to 60°C over 20 minutes, and subsequently from 60°C to 24°C over a period of 14 hours.

### Purification of NRs by PEG Precipitation

The purification process to separate the NRs from free staples was performed via polyethylene glycol (PEG) precipitation. The buffers prepared for this process included 2X PEG Buffer (10 mM Tris, 2 mM EDTA, and 1010 mM NaCl), PEG Buffer (15% PEG8000 in 1X PEG Buffer) and Storage Buffer (1X PBS, pH 7.4, and 10 mM MgCl_2_). After the NR folding process, the reaction mixture was transferred to a DNA LoBind tube (Eppendorf). An equal volume (1:1 v/v) of PEG buffer (15% PEG8000 in 1X PEG Buffer) was added, and the mixture was gently mixed. This mixture was incubated on ice for 10 minutes. To isolate the NRs, the reaction mixture was centrifuged at 21,000 x g for 25 minutes at 16°C. The supernatant was carefully removed, and the pellet was resuspended in 25 μL of storage buffer. The pellet was then incubated for 30 minutes at 30°C. Finally, the concentration of the NRs was determined by measuring the absorbance at 260 nm, using an extinction coefficient (ε) value of 94,500,000 M-^1^·cm-^1^.

### Oligo conjugation to antiEGFR-Antibody

To facilitate oligonucleotide conjugation to the antibody, a reaction mixture was prepared combining the antibody with 5 molar equivalents of NHS ester-(PEG)_4_-DBCO (dibenzocyclooctyne) and 3 molar equivalents of NHS ester Cy3 in a low protein binding tube. The linkers were first dissolved in DMSO and the antibody in 1X PBS was then added to the mixture. The reaction was carried out in a thermoshaker for 6 hours at room temperature at 300 RPM. Post-reaction, excess DBCO and Cy3 linkers were removed by overnight dialysis against 1x PBS using a dialysis membrane tubing with a 3 kDa molecular weight cutoff (MWCO) (SnakeSkin Dialysis Tubing, Thermofisher Scientific). Following dialysis, the sample was concentrated via ultracentrifugation using a filter with a 100 kDa MWCO (Merck Millipore). The concentration was determined by measuring absorbance at 280 nm (ε = 210,000 cm-^1^ M-^1^).

The obtained Cy3-/DBCO-Antibody was then reacted with 5 molar excess of an oligonucleotide modified with an azide group (N_3_). This reaction mixture was gently stirred and allowed to proceed overnight at 4°C. Subsequent purification of the product was conducted using a 50 Kda MWCO filter. The final results were visualized using SDS-PAGE gel electrophoresis.

### SDS-PAGE gel electrophoresis

Each sample contained 5 µg of protein dissolved in 1X PBS buffer. 2X loading dye containing DTT was added in a 1:1 ratio in each sample. Before loading, the samples were denatured at 95°C for 5 minutes, and 14 µL of each sample were loaded into the wells, alongside 5 µL of protein ladder (Precision Plus Protein, Bio-Rad). The gel electrophoresis was conducted in a pre-cast gel (Mini-Protean TGX 4-20% gel) in Tris/glycine buffer for 55 minutes at 150V.

### NR – Antibody/Aptamer conjugation

The conjugation of DNA-labeled antibodies or aptamers to purified DNA NRs was performed by incubating the NRs with the oligomodified anti-EGFR antibody/aptamer at a 3-fold molar excess relative to the handles. The anti-EGFR RNA aptamer was prepared as described previously in the work of Delcanale, P. *et al*.^46^ The labeling process was conducted for 1 hour at 37°C, followed by 2 hours at 22°C in 1X PBS (pH 7.4) containing 10 mM MgCl_2_. The results were analyzed by electrophoresis on a 1.5% agarose gel. In particular, a final concentration of 10 nM of NRs (empty or conjugated with ligands) were loaded in the wells. The run was performed in 0.5XTBE buffer for 90 minutes at 75V. To ensure the stability of the NRs, the run was cooled in an ice bath.

### Cell Culture

The MDA-MB-468 (ATCC cat. HTB-132) and Hek 293T (ATCC cat. CRL-3519) cell lines were cultured in Thermo Scientific™ Nunc™ Cell Culture Treated Flasks with Filter Caps. The cells were maintained in Dulbecco’s Modified Eagle Medium (DMEM, Thermofisher Scientific) supplemented with 5% fetal bovine serum (FBS, Thermofisher Scientific) and 1% Penicillin-Streptomycin (P/S, Thermofisher Scientific). Incubation was carried out at 37°C with 5% CO_2_.

### Confocal laser scanning microscopy

Approximately 10^4^ cells were seeded into a microscopy sample dish (µ-Slide 8 Well. Ibidi, no. #1.5) in a final volume of 300 µL. The slide was then placed in the incubator (5% CO2, 37°C) and cells were allowed to adhere overnight. Prior to imaging, the medium was exchanged with 1X Hanks’ Balanced Salt Solution (HBSS, Sigma-Aldrich) containing 20 nM of NRs. After 1 h of NP incubation, the cell membrane was stained with BioTracker™ 400 Blue Cytoplasmic Membrane Dye (Sigma Aldrich) according to the suppliers’ instructions. The cells and NRs were visualized using the Nikon STEDYCON microscope operating in confocal mode equipped with a 60x oil immersion objective.

### Single Particle Tracking Experiments

Approximately 10^4^ cells were seeded into a microscopy sample dish (µ-Slide 8 Well. Ibidi, no. #1.5) in a final volume of 300 µL. The slide was then placed in the incubator (5% CO2, 37°C) and cells were allowed to adhere overnight. Prior to imaging, the medium was exchanged with 1X Hanks’ Balanced Salt Solution (HBSS, Sigma-Aldrich) containing 1 nM of NRs. The NRs were visualized using the Oxford Nanoimager microscope in HiLo mode at an angle of 51°, exciting with a 635 nm laser operating at 25mW. The microscope was equipped with a 100x oil immersion objective. For each experimental condition, a total of 5 acquisitions were performed at three distinct time points: 10 minutes, 30 minutes, and 1 hour after the addition of the NR-containing solution to the cells. Each recorded movie comprised 2000 frames, with an exposure time of 10 ms for each frame (100 frames per second). Single-particle tracking was performed using the NimOS software, and the results were filtered with the following settings: maximum frame gap of 3, minimum distance between frames of 0.800 µm, exclusion radius of 1.200 µm, and a minimum number of steps of 30. With these parameters, the tracks and diffusion coefficient (D, μm^2^s-^1^) were extracted for each nanoparticle that met the applied filtering criteria. The analysis of motion (mean square displacement) was performed with the Python-based Nano-micromotor Analysis Tool (NMAT) v0.7 (https://github.com/rafamestre/NMAT-nanomicromotor-analysis-tool)^51^.

To measure kinetics parameters, only trajectories with a diffusion coefficient lower than 1 μm^2^s-^1^ and longer than 1 second (100 steps) were considered. This is done to discard free-floating NRs and non-specific short interactions, respectively. The relevant equations are described below:

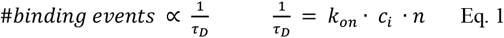

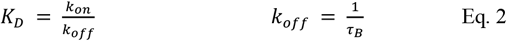

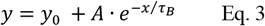

To obtain the value of τ_B_ (average dwell time of the interaction), the distribution of binding times were fitted with Eq.3 in OriginLab 2020 software.

## Supplementary information

### Supplementary tables

**Supplementary Table 1:**
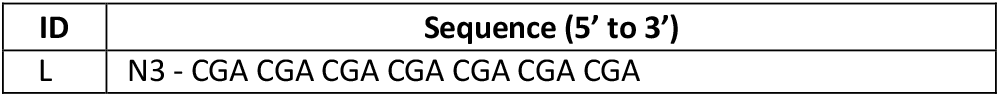
Anti-handle for Anti-EGFR Antibody/Aptamer labeling. This 3’azide-oligo was used for NHS coupling to the Anti-EGFR Antibody or Aptamer.

**Supplementary Table 2:**
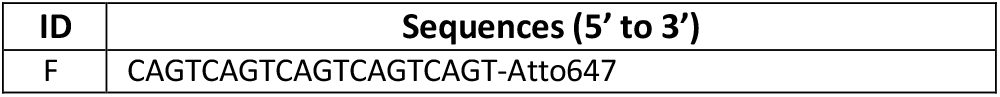
Fluorescently labeled imagers. To label DNA nanostructures, complementary imager (F) strands were used during the self-assembly of the DNA nanostructure.

**Supplementary Table 3:**
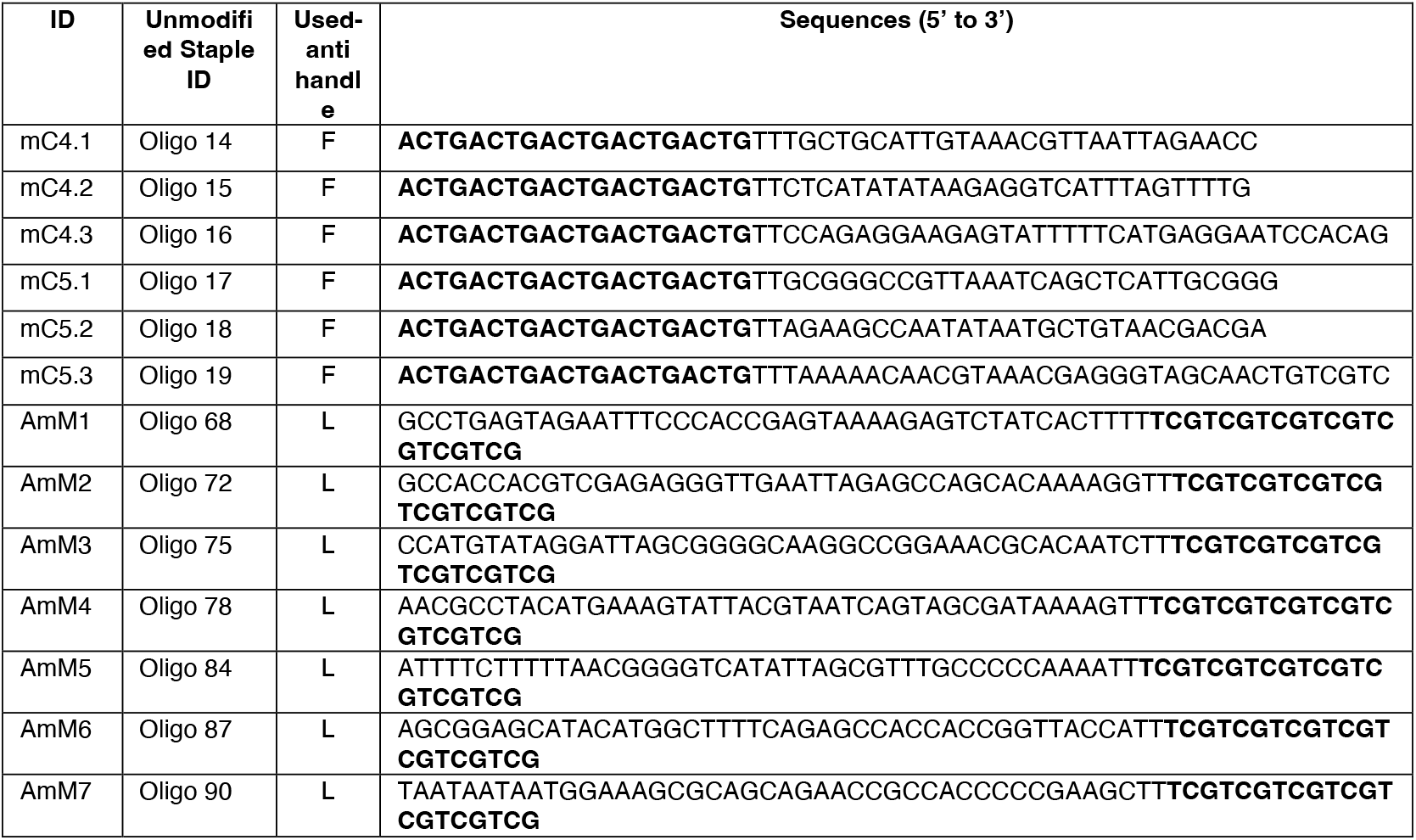
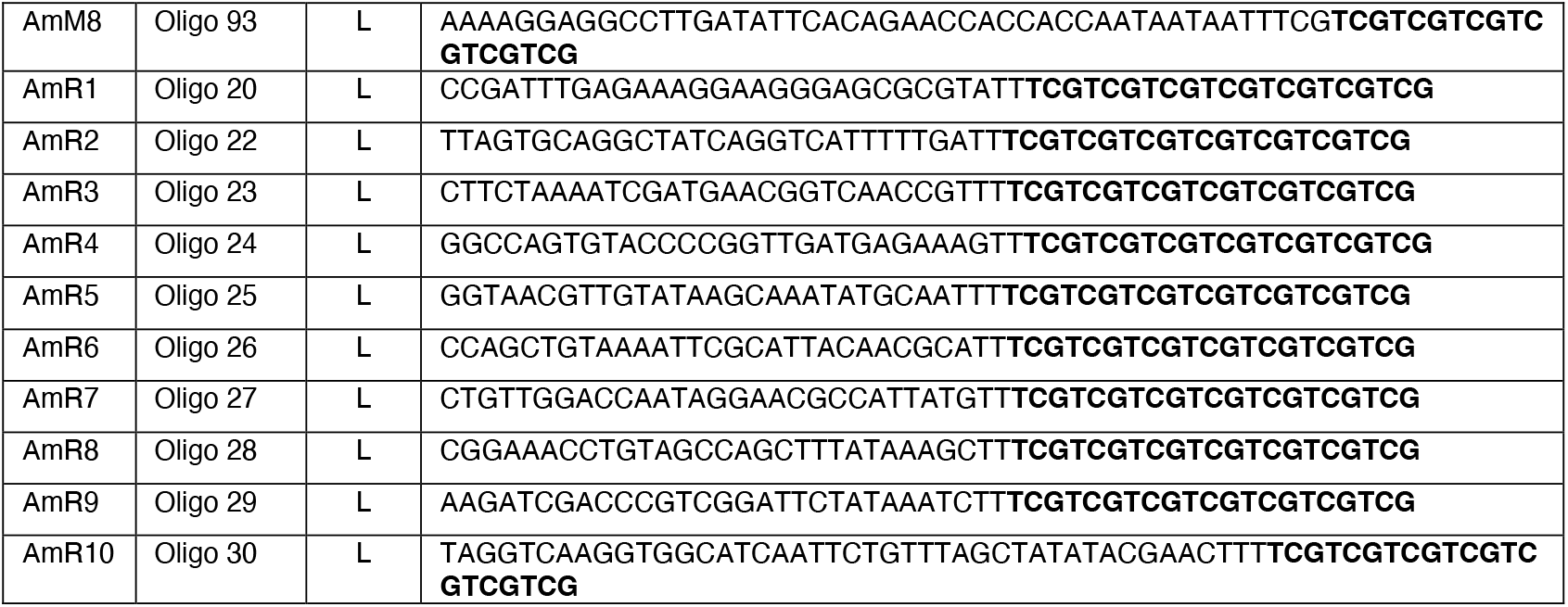
Handle-extended staple strands antibody incorporation. To assemble DNA nanorods for anti-EGFR Antibody/Aptamer, appropriate unmodified staples were replaced with handle-extended staple strands. Bold nucleotides refer to the extended handle sequence. Labels: A, antibody/aptamer; C, fluorophore.

**Supplementary Table 4:**
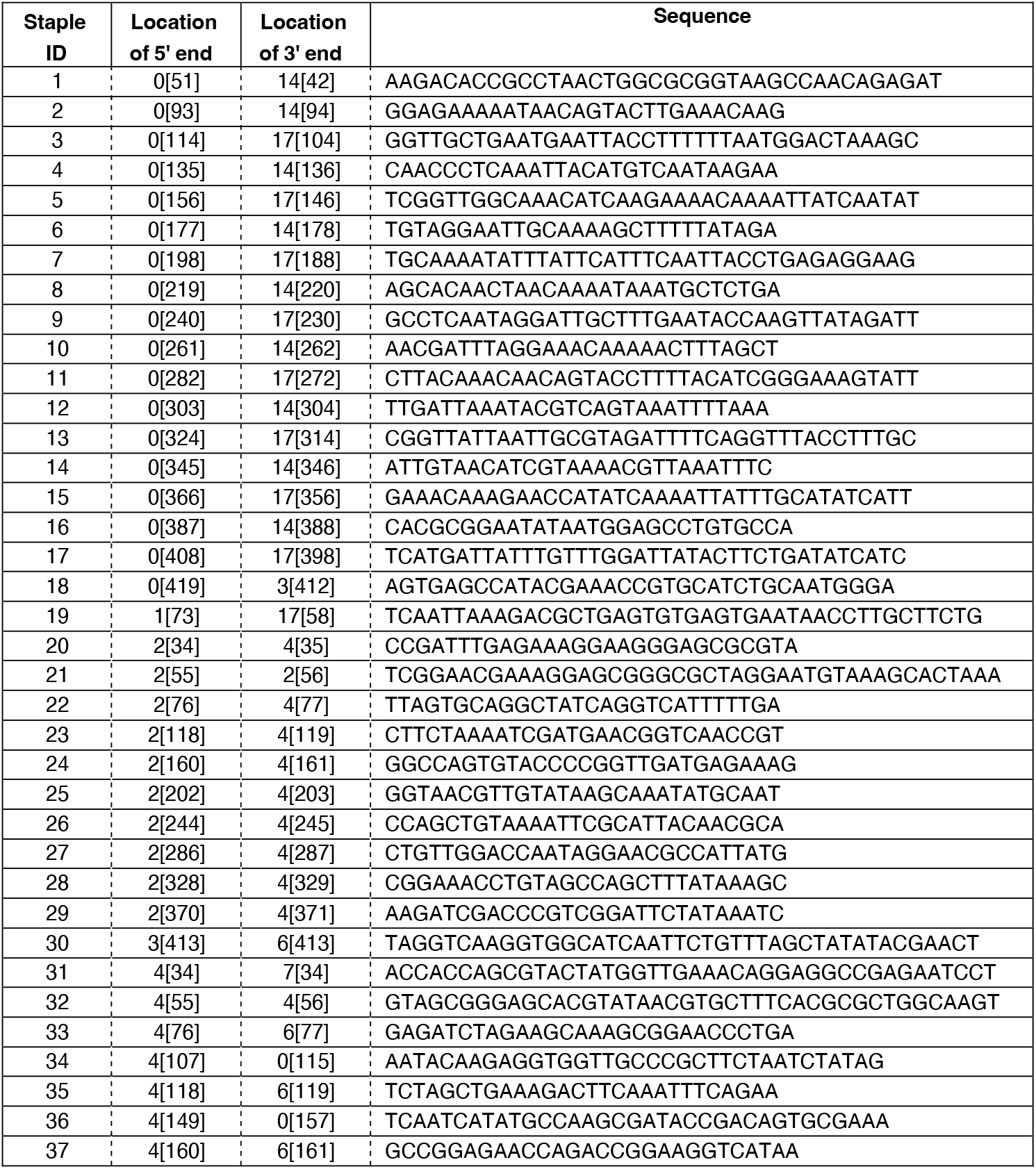
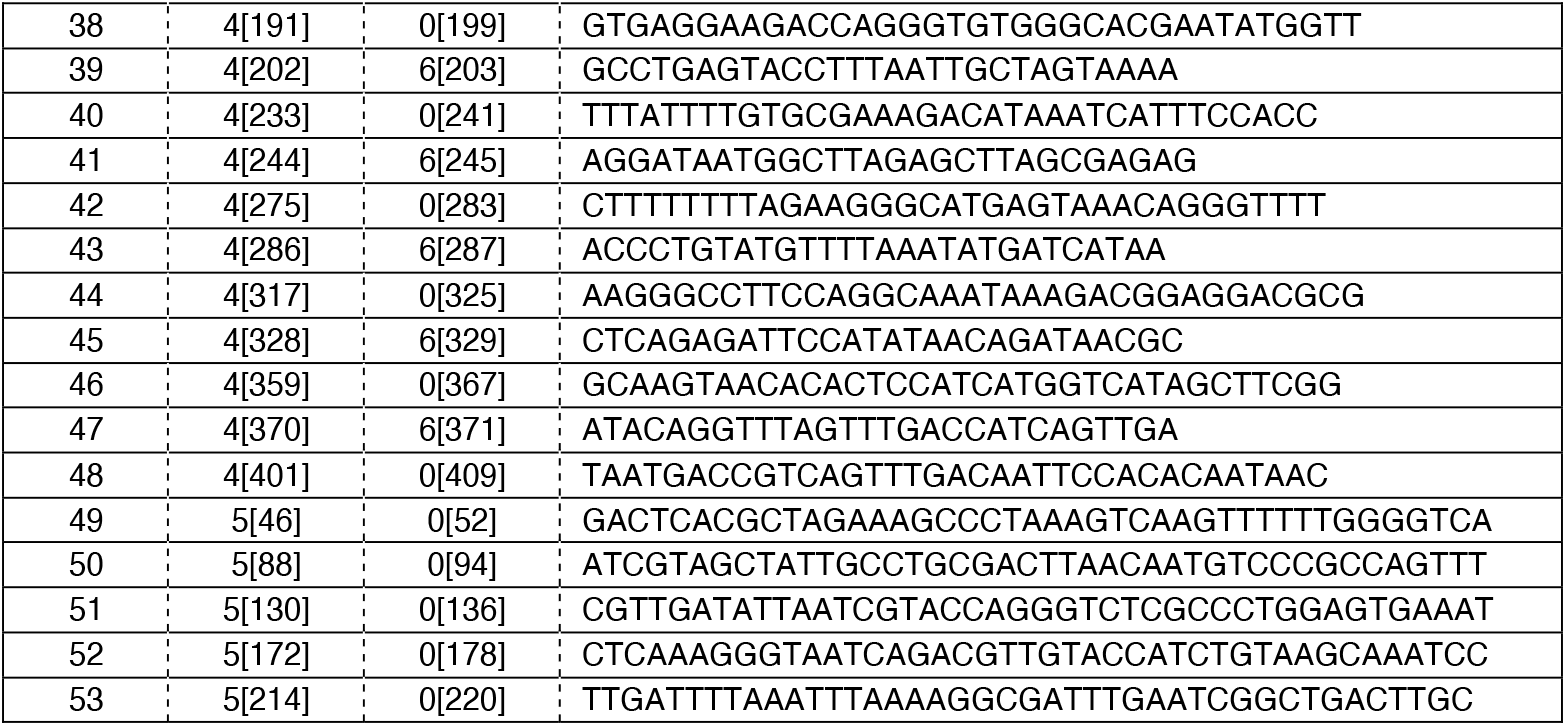
Sequences of unmodified staple strands of the DNA nanorod. Locations of the 5’ and 3’ end are indicated using the reference helix number, with the reference nucleotide position denoted in brackets.

### Supplementary figures

**Figure S1:**
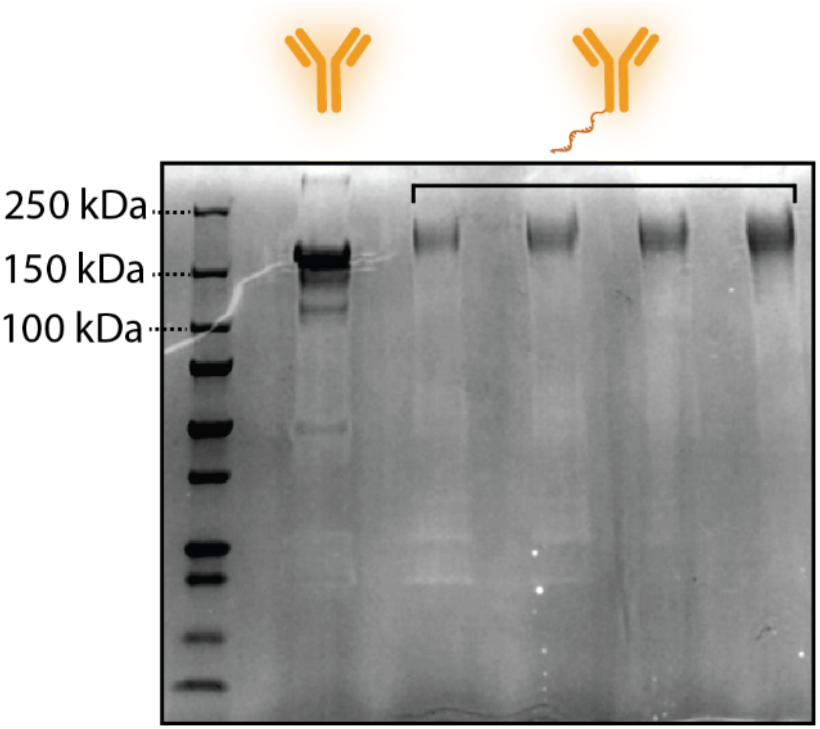
non-denaturing SDS page gel of the EGFR antibody (first lane) and the oligo conjugated EGFR antibody (lane 2-5, increasing concentration of antibody)

**Figure S2:**
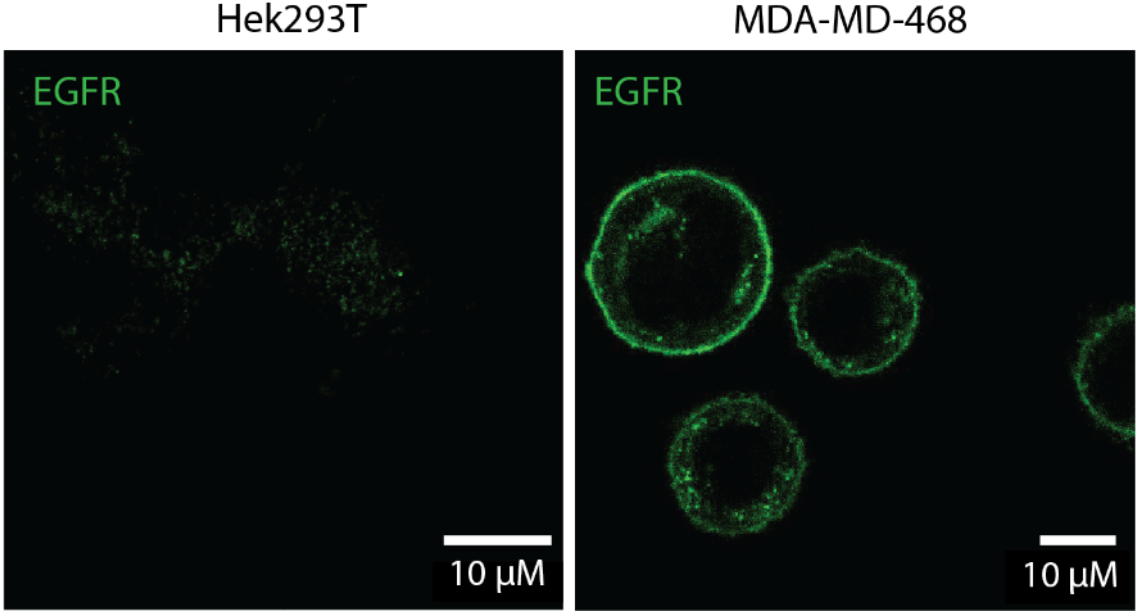
Immunostaining of the EGFR on Hek 293T cells (left) and the MDA MD 468 cells (right). Images were taken via confocal laser scanning microscopy.

**Figure S3:**
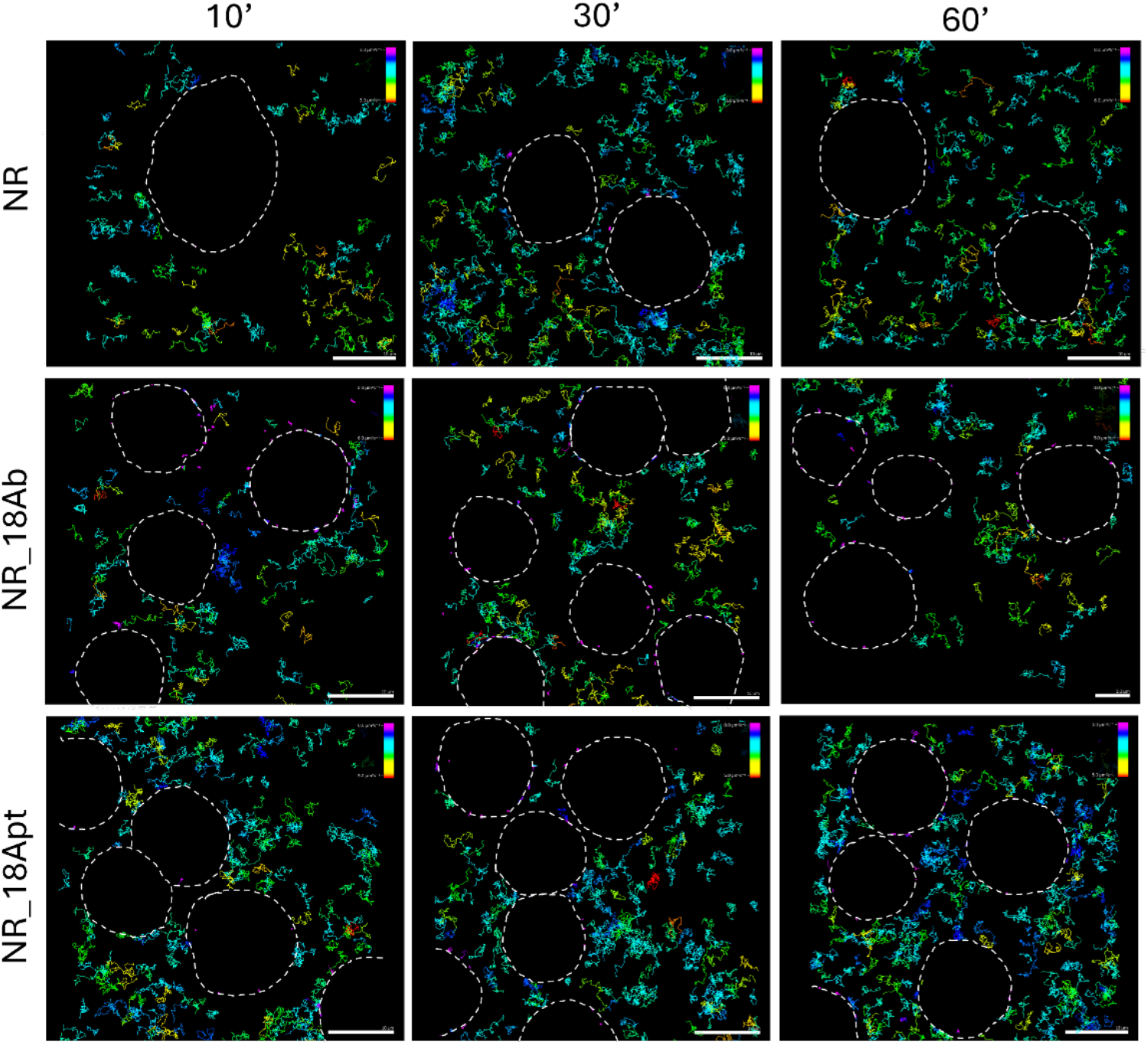
Plots of the trajectories of the different NR designs (empty NR, NR_18Ab and NR_18Apt) 10, 30 and 60 min after NR administration to MDA MD 468 cells. The cell contours are indicated by the dotted line. Scale bars values are displayed on the image, the color bar represents variations in diffusion coefficients (ranging from 0 to 6 µm2/s). Scale bar is 10 µm.

**Figure S4:**
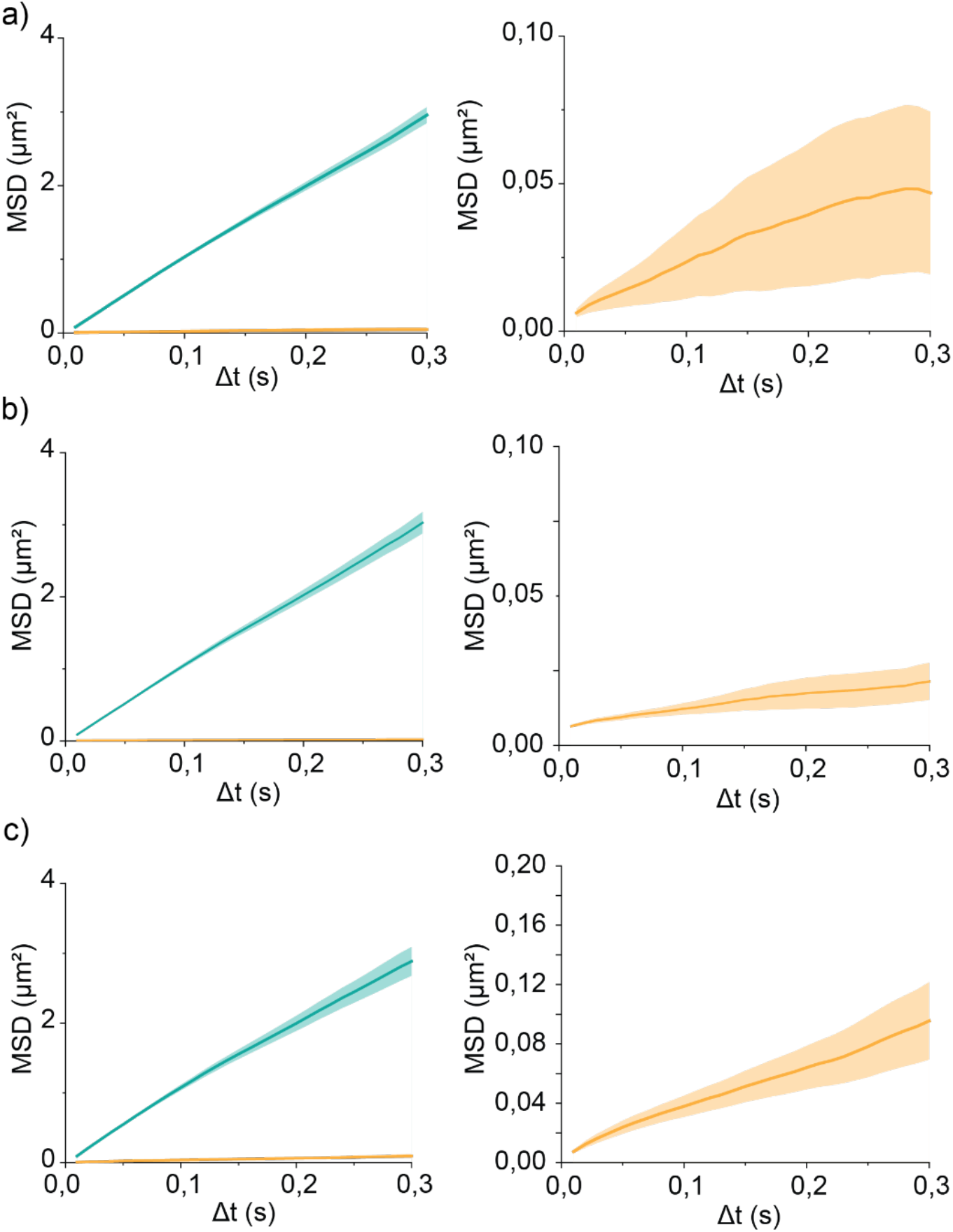
Plots of the mean squared displacement over time for a) the empty NR, b) NR_18Ab and c) NR_18Apt. Trajectories that have a diffusion coefficient higher than 1 (free diffusing NRs) are plotted in blue whereas trajectories with a diffusion coefficient lower or equal to 1 (receptor binding NRs) are plotted in yellow. Since the receptor binding NRs have a very low mobility, the yellow line is located too close to the x-axis. Therefore, the axes were rescaled to clarify the trend of the receptor binding NR population (second column).

**Figure S5:**
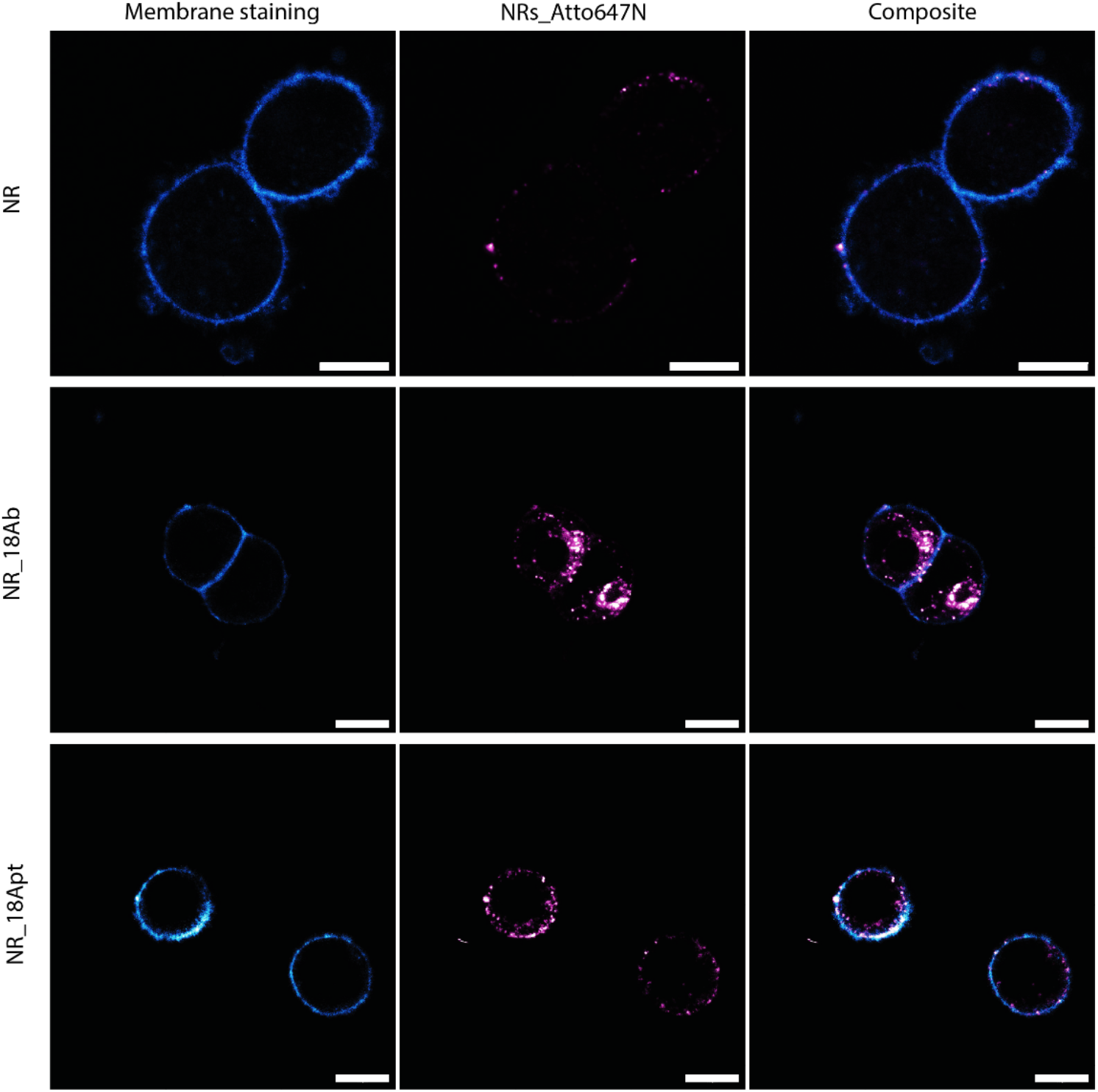
Confocal laser scanning microscopy images of MDA MD 468 cells incubated with empty NRs, NRs_18Ab and NR_18Apt for 24 hours, all labeled with Atto648N (magenta). The membrane is stained with BioTracker™ 400 Blue Cytoplasmic Membrane Dye (cyan). Scale bar is 10 µm.

**Figure S6:**
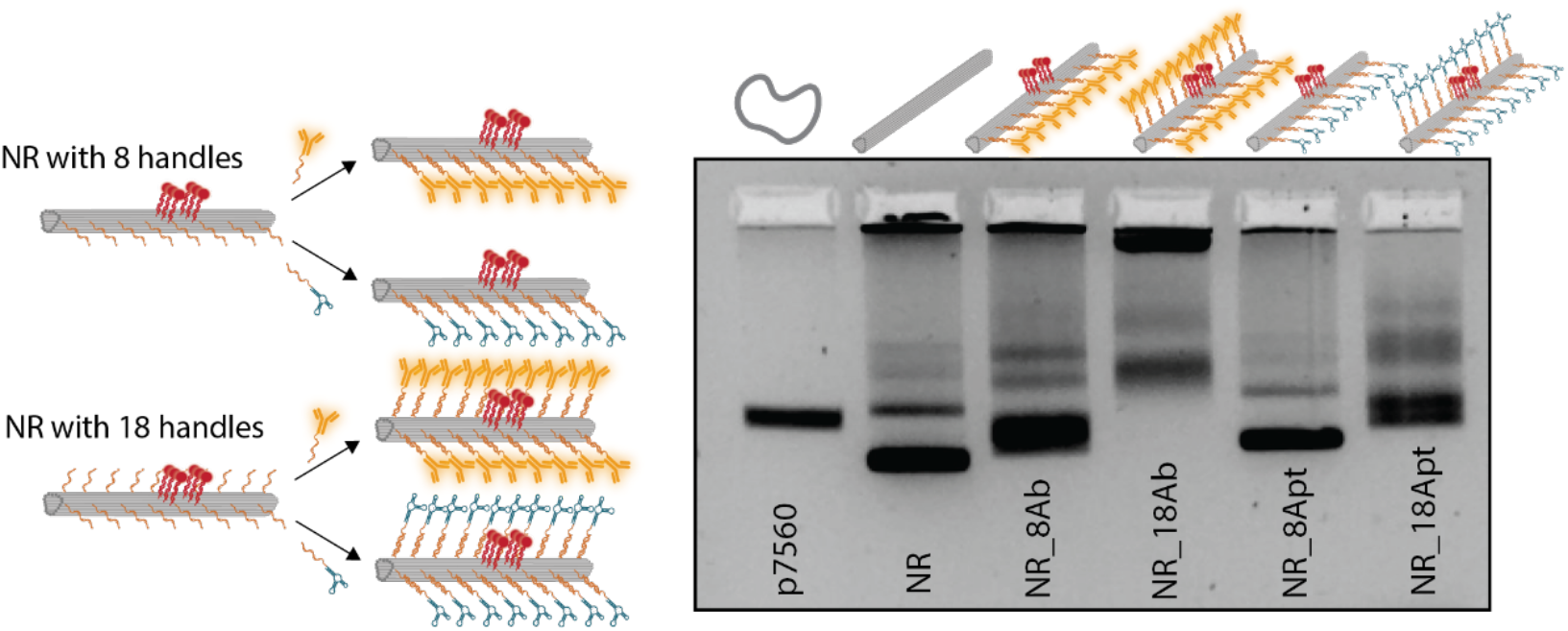
Agarose gel of nanorods conjugated with 8 or 18 EGFR antibodies and 8 or 18 aptamers.

## References

(1) Manz, B. N.; Groves, J. T. Spatial Organization and Signal Transduction at Intercellular Junctions. Nat Rev Mol Cell Biol 2010, 11 (5), 342–352. 10.1038/nrm2883.

(2) Waldman, A. D.; Fritz, J. M.; Lenardo, M. J. A Guide to Cancer Immunotherapy: From T Cell Basic Science to Clinical Practice. Nat Rev Immunol 2020, 20 (11), 651–668. 10.1038/s41577-020-0306-5.

(3) Huppa, J. B.; Davis, M. M. T-Cell-Antigen Recognition and the Immunological Synapse. Nat Rev Immunol 2003, 3 (12), 973–983. 10.1038/nri1245.

(4) Winkler, J.; Abisoye-Ogunniyan, A.; Metcalf, K. J.; Werb, Z. Concepts of Extracellular Matrix Remodelling in Tumour Progression and Metastasis. Nat Commun 2020, 11 (1), 5120. 10.1038/s41467-020-18794-x.

(5) Singer, S. J. Intercellular Communication and Cell-Cell Adhesion. Science 1992, 255 (5052), 1671–1677. 10.1126/science.1313187.

(6) Motsch, V.; Brameshuber, M.; Baumgart, F.; Schütz, G. J.; Sevcsik, E. A Micropatterning Platform for Quantifying Interaction Kinetics between the T Cell Receptor and an Intracellular Binding Protein. Sci Rep 2019, 9 (1), 3288. 10.1038/s41598-019-39865-0.

(7) Huang, J.; Zarnitsyna, V. I.; Liu, B.; Edwards, L. J.; Jiang, N.; Evavold, B. D.; Zhu, C. The Kinetics of Two-Dimensional TCR and pMHC Interactions Determine T-Cell Responsiveness. Nature 2010, 464 (7290), 932–936. 10.1038/nature08944.

(8) Choi, Y.; Cho, B. K.; Seok, S. H.; Kim, C.; Ryu, J. H.; Kwon, I. C. Controlled Spatial Characteristics of Ligands on Nanoparticles: Determinant of Cellular Functions. Journal of Controlled Release 2023, 360, 672–686. 10.1016/j.jconrel.2023.07.020.

(9) Cifuentes-Rius, A.; Desai, A.; Yuen, D.; Johnston, A. P. R.; Voelcker, N. H. Inducing Immune Tolerance with Dendritic Cell-Targeting Nanomedicines. Nat. Nanotechnol. 2021, 16 (1), 37–46. 10.1038/s41565-020-00810-2.

(10) Woythe, L.; Tito, N. B.; Albertazzi, L. A Quantitative View on Multivalent Nanomedicine Targeting. Advanced Drug Delivery Reviews 2021, 169, 1–21. 10.1016/j.addr.2020.11.010.

(11) Rothemund, P. W. K. Folding DNA to Create Nanoscale Shapes and Patterns. Nature 2006, 440 (7082), 297–302. 10.1038/nature04586.

(12) Chen, J. H.; Seeman, N. C. Synthesis from DNA of a Molecule with the Connectivity of a Cube. Nature 1991, 350 (6319), 631–633. 10.1038/350631a0.

(13) Keller, A.; Linko, V. Challenges and Perspectives of DNA Nanostructures in Biomedicine. Angew Chem Int Ed Engl 2020, 59 (37), 15818–15833. 10.1002/anie.201916390.

(14) Wei, B.; Dai, M.; Yin, P. Complex Shapes Self-Assembled from Single-Stranded DNA Tiles. Nature 2012, 485 (7400), 623–626. 10.1038/nature11075.

(15) Veneziano, R.; Ratanalert, S.; Zhang, K.; Zhang, F.; Yan, H.; Chiu, W.; Bathe, M. Designer Nanoscale DNA Assemblies Programmed from the Top Down. Science 2016, 352 (6293), 1534. 10.1126/science.aaf4388.

(16) Douglas, S. M.; Marblestone, A. H.; Teerapittayanon, S.; Vazquez, A.; Church, G. M.; Shih, W. M. Rapid Prototyping of 3D DNA-Origami Shapes with caDNAno. Nucleic Acids Research 2009, 37 (15), 5001–5006. 10.1093/nar/gkp436.

(17) Seeman, N. C. DNA Nanotechnology: Novel DNA Constructions. Annu Rev Biophys Biomol Struct 1998, 27, 225–248. 10.1146/annurev.biophys.27.1.225.

(18) Seeman, N. C.; Sleiman, H. F. DNA Nanotechnology. Nat Rev Mater 2017, 3 (1), 1–23. 10.1038/natrevmats.2017.68.

(19) Rinker, S.; Ke, Y.; Liu, Y.; Chhabra, R.; Yan, H. Self-Assembled DNA Nanostructures for Distance-Dependent Multivalent Ligand–Protein Binding. Nature Nanotech 2008, 3 (7), 418–422. 10.1038/nnano.2008.164.

(20) Shaw, A.; Hoffecker, I. T.; Smyrlaki, I.; Rosa, J.; Grevys, A.; Bratlie, D.; Sandlie, I.; Michaelsen, T. E.; Andersen, J. T.; Högberg, B. Binding to Nanopatterned Antigens Is Dominated by the Spatial Tolerance of Antibodies. Nature Nanotech 2019, 14 (2), 184–190. 10.1038/s41565-018-0336-3.

(21) Kurisinkal, E. E.; Caroprese, V.; Koga, M. M.; Morzy, D.; Bastings, M. M. C. Selective Integrin A5β1 Targeting through Spatially Constrained Multivalent DNA-Based Nanoparticles. Molecules 2022, 27 (15), 4968. 10.3390/molecules27154968.

(22) Kwon, P. S.; Ren, S.; Kwon, S.-J.; Kizer, M. E.; Kuo, L.; Xie, M.; Zhu, D.; Zhou, F.; Zhang, F.; Kim, D.; Fraser, K.; Kramer, L. D.; Seeman, N. C.; Dordick, J. S.; Linhardt, R. J.; Chao, J.; Wang, X. Designer DNA Architecture Offers Precise and Multivalent Spatial Pattern-Recognition for Viral Sensing and Inhibition. Nat. Chem. 2020, 12 (1), 26–35. 10.1038/s41557-019-0369-8.

(23) Cremers, G. A. O.; Rosier, B. J. H. M.; Meijs, A.; Tito, N. B.; van Duijnhoven, S. M. J.; van Eenennaam, H.; Albertazzi, L.; de Greef, T. F. A. Determinants of Ligand-Functionalized DNA Nanostructure–Cell Interactions. J. Am. Chem. Soc. 2021, 143 (27), 10131–10142. 10.1021/jacs.1c02298.

(24) Comberlato, A.; Koga, M. M.; Nüssing, S.; Parish, I. A.; Bastings, M. M. C. Spatially Controlled Activation of Toll-like Receptor 9 with DNA-Based Nanomaterials. Nano Lett. 2022, 22 (6), 2506–2513. 10.1021/acs.nanolett.2c00275.

(25) Lee, D. S.; Qian, H.; Tay, C. Y.; Leong, D. T. Cellular Processing and Destinies of Artificial DNA Nanostructures. Chem. Soc. Rev. 2016, 45 (15), 4199–4225. 10.1039/C5CS00700C.

(26) Vinther, M.; Kjems, J. Interfacing DNA Nanodevices with Biology: Challenges, Solutions and Perspectives. New J. Phys. 2016, 18 (8), 085005. 10.1088/1367-2630/18/8/085005.

(27) Weiden, J.; Bastings, M. M. C. DNA Origami Nanostructures for Controlled Therapeutic Drug Delivery. Current Opinion in Colloid & Interface Science 2021, 52, 101411. 10.1016/j.cocis.2020.101411.

(28) Bila, H.; Kurisinkal, E. E.; Bastings, M. M. C. Engineering a Stable Future for DNA-Origami as a Biomaterial. Biomater. Sci. 2019, 7 (2), 532–541. 10.1039/C8BM01249K.

(29) Mei, Q.; Wei, X.; Su, F.; Liu, Y.; Youngbull, C.; Johnson, R.; Lindsay, S.; Yan, H.; Meldrum, D. Stability of DNA Origami Nanoarrays in Cell Lysate. Nano Lett 2011, 11 (4), 1477–1482. 10.1021/nl1040836.

(30) Linko, V.; Keller, A. Stability of DNA Origami Nanostructures in Physiological Media: The Role of Molecular Interactions. Small 2023, 19 (34), 2301935. 10.1002/smll.202301935.

(31) Zhao, Y.-X.; Shaw, A.; Zeng, X.; Benson, E.; Nyström, A. M.; Högberg, B. DNA Origami Delivery System for Cancer Therapy with Tunable Release Properties. ACS Nano 2012, 6 (10), 8684–8691. 10.1021/nn3022662.

(32) Seitz, I.; Ijäs, H.; Linko, V.; Kostiainen, M. A. Optically Responsive Protein Coating of DNA Origami for Triggered Antigen Targeting. ACS Appl. Mater. Interfaces 2022, 14 (34), 38515–38524. 10.1021/acsami.2c10058.

(33) Pal, S.; Rakshit, T. Folate-Functionalized DNA Origami for Targeted Delivery of Doxorubicin to Triple-Negative Breast Cancer. Front. Chem. 2021, 9. 10.3389/fchem.2021.721105.

(34) Wang, Z.; Song, L.; Liu, Q.; Tian, R.; Shang, Y.; Liu, F.; Liu, S.; Zhao, S.; Han, Z.; Sun, J.; Jiang, Q.; Ding, B. A Tubular DNA Nanodevice as a siRNA/Chemo-Drug Co-Delivery Vehicle for Combined Cancer Therapy. Angew Chem Int Ed Engl 2021, 60 (5), 2594–2598. 10.1002/anie.202009842.

(35) Pan, Q.; Nie, C.; Hu, Y.; Yi, J.; Liu, C.; Zhang, J.; He, M.; He, M.; Chen, T.; Chu, X. Aptamer-Functionalized DNA Origami for Targeted Codelivery of Antisense Oligonucleotides and Doxorubicin to Enhance Therapy in Drug-Resistant Cancer Cells. ACS Appl. Mater. Interfaces 2020, 12 (1), 400–409. 10.1021/acsami.9b20707.

(36) Ge, Z.; Guo, L.; Wu, G.; Li, J.; Sun, Y.; Hou, Y.; Shi, J.; Song, S.; Wang, L.; Fan, C.; Lu, H.; Li, Q. DNA Origami-Enabled Engineering of Ligand–Drug Conjugates for Targeted Drug Delivery. Small 2020, 16 (16), 1904857. 10.1002/smll.201904857.

(37) Lee, H.; Lytton-Jean, A. K. R.; Chen, Y.; Love, K. T.; Park, A. I.; Karagiannis, E. D.; Sehgal, A.; Querbes, W.; Zurenko, C. S.; Jayaraman, M.; Peng, C. G.; Charisse, K.; Borodovsky, A.; Manoharan, M.; Donahoe, J. S.; Truelove, J.; Nahrendorf, M.; Langer, R.; Anderson, D. G. Molecularly Self-Assembled Nucleic Acid Nanoparticles for Targeted in Vivo siRNA Delivery. Nature Nanotech 2012, 7 (6), 389–393. 10.1038/nnano.2012.73.

(38) Wu, X.; Yang, C.; Wang, H.; Lu, X.; Shang, Y.; Liu, Q.; Fan, J.; Liu, J.; Ding, B. Genetically Encoded DNA Origami for Gene Therapy In Vivo. J. Am. Chem. Soc. 2023, 145 (16), 9343–9353. 10.1021/jacs.3c02756.

(39) Li, S.; Jiang, Q.; Liu, S.; Zhang, Y.; Tian, Y.; Song, C.; Wang, J.; Zou, Y.; Anderson, G. J.; Han, J. Y.; Chang, Y.; Liu, Y.; Zhang, C.; Chen, L.; Zhou, G.; Nie, G.; Yan, H.; Ding, B.; Zhao, Y. A DNA Nanorobot Functions as a Cancer Therapeutic in Response to a Molecular Trigger in Vivo. Nature biotechnology 2018, 36 (3), 258–264. 10.1038/nbt.4071.

(40) Wagenbauer, K. F.; Pham, N.; Gottschlich, A.; Kick, B.; Kozina, V.; Frank, C.; Trninic, D.; Stömmer, P.; Grünmeier, R.; Carlini, E.; Tsiverioti, C. A.; Kobold, S.; Funke, J. J.; Dietz, H. Programmable Multispecific DNA-Origami-Based T-Cell Engagers. Nat. Nanotechnol. 2023, 18 (11), 1319–1326. 10.1038/s41565-023-01471-7.

(41) Xu, T.; Yu, S.; Sun, Y.; Wu, S.; Gao, D.; Wang, M.; Wang, Z.; Tian, Y.; Min, Q.; Zhu, J.-J. DNA Origami Frameworks Enabled Self-Protective siRNA Delivery for Dual Enhancement of Chemo-Photothermal Combination Therapy. Small 2021, 17 (46), 2101780. 10.1002/smll.202101780.

(42) Zeng, Y. C.; Young, O. J.; Wintersinger, C. M.; Anastassacos, F. M.; MacDonald, J. I.; Isinelli, G.; Dellacherie, M. O.; Sobral, M.; Bai, H.; Graveline, A. R.; Vernet, A.; Sanchez, M.; Mulligan, K.; Choi, Y.; Ferrante, T. C.; Keskin, D. B.; Fell, G. G.; Neuberg, D.; Wu, C. J.; Mooney, D. J.; Kwon, I. C.; Ryu, J. H.; Shih, W. M. Fine Tuning of CpG Spatial Distribution with DNA Origami for Improved Cancer Vaccination. Nat. Nanotechnol. 2024, 1–11. 10.1038/s41565-024-01615-3.

(43) Woythe, L.; Madhikar, P.; Feiner-Gracia, N.; Storm, C.; Albertazzi, L. A Single-Molecule View at Nanoparticle Targeting Selectivity: Correlating Ligand Functionality and Cell Receptor Density. ACS Nano 2022, 16 (3), 3785–3796. 10.1021/acsnano.1c08277.

(44) Guerrab, A. E.; Bamdad, M.; Kwiatkowski, F.; Bignon, Y.-J.; Penault-Llorca, F.; Aubel, C. Anti-EGFR Monoclonal Antibodies and EGFR Tyrosine Kinase Inhibitors as Combination Therapy for Triple-Negative Breast Cancer. Oncotarget 2016, 7 (45), 73618–73637. 10.18632/oncotarget.12037.

(45) Zhang, F.; Wang, S.; Yin, L.; Yang, Y.; Guan, Y.; Wang, W.; Xu, H.; Tao, N. Quantification of Epidermal Growth Factor Receptor Expression Level and Binding Kinetics on Cell Surfaces by Surface Plasmon Resonance Imaging. Anal Chem 2015, 87 (19), 9960–9965. 10.1021/acs.analchem.5b02572.

(46) Delcanale, P.; Porciani, D.; Pujals, S.; Jurkevich, A.; Chetrusca, A.; Tawiah, K. D.; Burke, D. H.; Albertazzi, L. Aptamers with Tunable Affinity Enable Single-Molecule Tracking and Localization of Membrane Receptors on Living Cancer Cells. Angew Chem Int Ed Engl 2020, 59 (42), 18546–18555. 10.1002/anie.202004764.

(47) Martinez-Veracoechea, F. J.; Frenkel, D. Designing Super Selectivity in Multivalent Nano-Particle Binding. 2011. 10.1073/pnas.1105351108.

(48) Curk, T.; Dobnikar, J.; Frenkel, D. Optimal Multivalent Targeting of Membranes with Many Distinct Receptors. 2017. 10.1073/pnas.1704226114.

(49) Hoffecker, I. T.; Shaw, A.; Sorokina, V.; Smyrlaki, I.; Högberg, B. The Geometric Determinants of Programmed Antibody Migration and Binding on Multi-Antigen Substrates. bioRxiv October 12, 2020, p 2020.10.12.336164. 10.1101/2020.10.12.336164.

(50) Veneziano, R.; Moyer, T. J.; Stone, M. B.; Wamhoff, E.-C.; Read, B. J.; Mukherjee, S.; Shepherd, T. R.; Das, J.; Schief, W. R.; Irvine, D. J.; Bathe, M. Role of Nanoscale Antigen Organization on B-Cell Activation Probed Using DNA Origami. Nat. Nanotechnol. 2020, 15 (8), 716–723. 10.1038/s41565-020-0719-0.

(51) Mestre, R.; Palacios, L. S.; Miguel-López, A.; Arqué, X.; Pagonabarraga, I.; Sánchez, S. Extraction of the Propulsive Speed of Catalytic Nano- and Micro-Motors under Different Motion Dynamics. arXiv July 30, 2020. 10.48550/arXiv.2007.15316.

